# High-Precision Pneumatic Induction of Traumatic Brain Injury in Larval Zebrafish

**DOI:** 10.64898/2026.02.24.707859

**Authors:** Kunrong Wang, Ping Zhang, Yijie Geng

## Abstract

Zebrafish have recently emerged as a cost-effective vertebrate model for Traumatic brain injury (TBI) research. However, current injury paradigms, particularly the weight-drop method, suffer from poor reproducibility due to sensitivity to small variations in materials and setup. Here, we introduce a high-precision pneumatic piston system, the Zebrafish Pneumatic Injury Device (ZePID), which delivers controlled, repeatable pressure pulses to induce TBI in larval zebrafish. ZePID achieves 97% accuracy in applied force and substantially reduces variability compared with weight-drop approaches. Using 6-days-post-fertilization (dpf) larvae, we quantified injury severity by tracking seizure-like locomotor behaviors. Larvae exposed to 150 psi exhibited a significant increase in maximum swimming speed relative to uninjured controls, as well as increased total distance traveled, consistent with TBI-associated hyperactivity. Overall, ZePID provides a standardized, compact, and high-efficiency method for inducing TBI in larval zebrafish.

## INTRODUCTION

Traumatic brain injury (TBI) is a major global public health concern and one of the leading causes of death and disability across all age groups. Each year, more than 69 million individuals worldwide experience TBI, with mild cases comprising approximately 80% of all incidents^1^. The condition arises from external mechanical forces such as blows, jolts, or penetrating injuries that disrupt normal brain function and may result in temporary or permanent cognitive, physical, and psychosocial impairments^2-4^. Epidemiologically, TBI affects diverse populations but is particularly prevalent among young adults, the elderly, and individuals in high-risk occupations or environments^5,6^. Pathophysiologically, TBI initiates a cascade of primary and secondary injuries involving excitotoxicity, oxidative stress, inflammation, and blood–brain barrier disruption, which collectively contribute to neuronal damage and long-term neurological deficits^7-10^. Despite advances in acute care, no definitive therapy exists to prevent secondary injury or restore full neurological function^11-14^.

Animal models remain essential for elucidating TBI pathophysiology and identifying candidate therapeutics. Because ethical and practical limitations restrict direct human experimentation, preclinical models provide critical insight into the complex cascade of primary and secondary injury mechanisms ranging from excitotoxicity and neuroinflammation to oxidative stress and apoptosis that occur after TBI^15^. Rodent models of TBI, including controlled cortical impact and fluid percussion paradigms, have been instrumental in elucidating cellular and molecular responses to injury^16-18^. Despite challenges in translating preclinical success to clinical efficacy, animal studies continue to generate essential data on neuroprotective targets and biomarkers that guide therapeutic discovery^19^. Moreover, innovative refinements in animal modeling, such as the use of hybrid models and genetically engineered species, are helping to bridge the translational gap^20^.

One widely used experimental paradigm for inducing diffuse TBI in rodents is the weight-drop model, in which a free-falling mass is guided through a vertical tube to deliver kinetic energy to the animal’s head. This impact produces acceleration and deformation of brain tissue without the need for craniotomy, resulting in widespread axonal and vascular injury that closely mimics aspects of human diffuse TBI^16^. Originally adapted from Marmarou’s model of diffuse axonal injury^21^, this technique has been extensively refined and validated across species and severities, demonstrating consistent histopathological, neuroinflammatory, and behavioral outcomes in rodent models^16,22-25^.

More recently, zebrafish have emerged as an attractive vertebrate system for modeling TBI^26,27^ due to their genetic, brain anatomical, and neurophysiological conservations with humans^28,29^, low maintenance cost, optical transparency, and regenerative capacity^30^. The development of a larval zebrafish weight-drop protocol^31,32^ demonstrated that this method can be adapted to zebrafish and can elicit measurable seizure-like responses. However, the zebrafish weight-drop paradigm exhibits several limitations. The system requires a long guide tube and precise alignment, and its performance is highly sensitive to small variations in tube geometry, friction, and environmental factors. These constraints can lead to poor reproducibility, variable force delivery, and low throughput, making it difficult to generate consistent injury severities or to scale the model for drug screening.

To address these challenges, inspired by methods recently developed in mouse models^33,34^, we developed a pneumatic piston–based TBI induction device that delivers controlled pressure pulses to larval zebrafish housed within a syringe chamber. By replacing the free-fall mass with electronically actuated pneumatic force, this system standardizes injury magnitude, improves reproducibility, and reduces the physical footprint of the apparatus. Establishing a more accurate and consistent TBI delivery method will strengthen the utility of zebrafish as a neurotrauma model and enhance their application in mechanistic studies and therapeutic screening.

## RESULTS

### Design and construction of the Zebrafish Pneumatic Injury Device (ZePID)

To overcome the variability and operator-dependent errors associated with the traditional weight-drop method, we developed a compact pneumatic system that replaces the free-falling mass and guide tube with electronically controlled pressure delivery. By eliminating the need for a one-meter guide tube, the resulting device reduces the overall footprint from 45 × 40 × 160 cm (Figure 1A) to 45 × 40 × 50 cm (Figure 1B), making it more practical for routine laboratory use.

**Figure 1.**
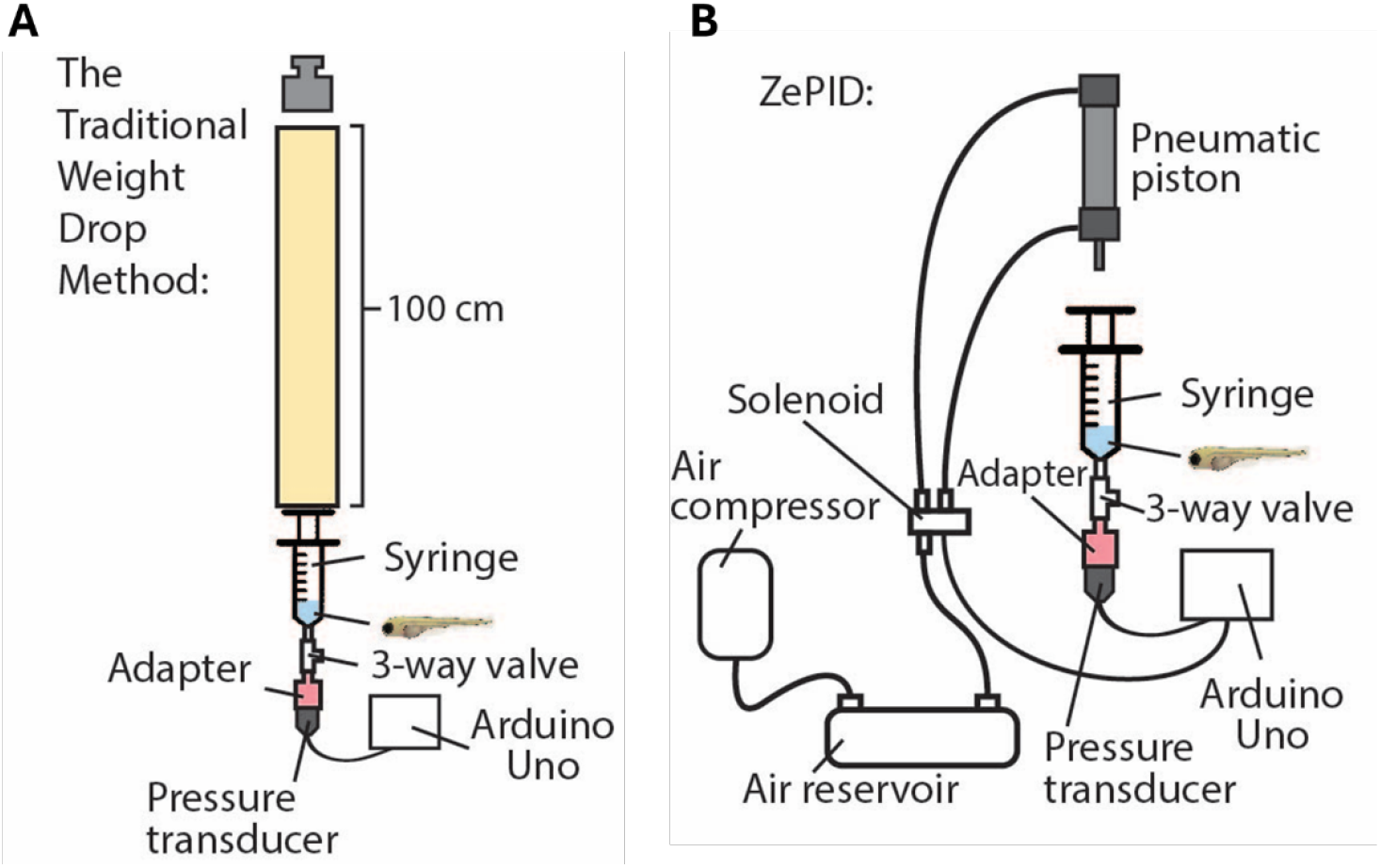
Comparing the traditional original weight drop method with the ZePID design. (**A**) A schematic demonstration of the traditional weight drop method. (**B**) A schematic demonstration of the ZePID system. The ZePID method improves on the original weight drop method by removing the guide tube and weight, replacing it with a pneumatic system controlled electrically by an Arduino microcontroller, which significantly improves accuracy and consistency. Pneumatic piston is triggered by the opening of the solenoid. Zebrafish larvae are confined in the chamber of a 20mL syringe. Pressure transducer attached to the device measures the pressure within the syringe at all times and verify the effectiveness of the pneumatic system.

The core of the system is a pneumatic piston (Baomain CDJ2B10-10-B) driven by compressed air supplied by a reservoir tank (Vex Robotics). A solenoid valve (SMC SY3120-6LZD-M5), connected to the piston via 4-mm pneumatic tubing, regulates the flow of air from the reservoir. The solenoid is controlled by an Arduino Uno Rev3 microcontroller, which triggers piston extension when activated. Upon firing, pressurized air is rapidly released through the solenoid into the piston chamber, producing an immediate, reproducible mechanical pulse at the programmed reservoir pressure (Figure 1B).

Larval zebrafish were housed during testing in a 20-mL syringe chamber connected to a pressure transducer (AUTEX 702373916802) that continuously recorded the pressure experienced by the fish. The syringe tip is connected to a 3-way Luer-lock valve through a custom 3D-printed adapter designed in Fusion 360 (Figure 2A & Supplementary Figure 1A). The adapter incorporates a hot-glued female Luer tip for attachment to the valve and threaded ports on the opposite side for integration with the pressure transducer (Figure 2B). This design allows the device to be opened easily at the valve–adapter junction when loading larvae, while maintaining a sealed and stable connection to the transducer during operation (Figure 2C).

**Figure 2.**
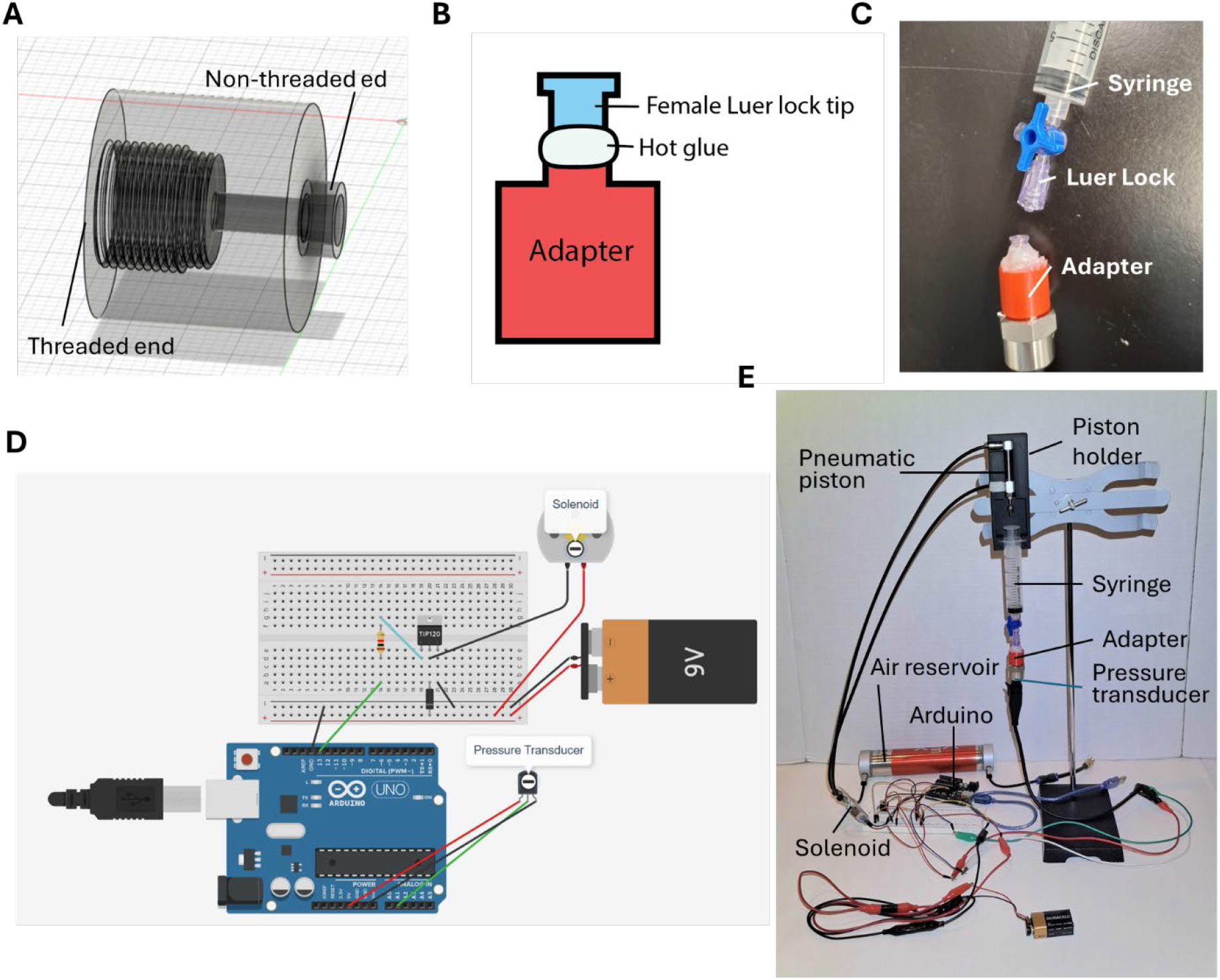
The construction of ZePID. (**A**) Isometric view of the adapter shown in Fusion 360, with the threaded end used to attach the pressure transducer and the non-threaded end joined with a female Luer lock tip. (**B**) A diagram showing the adapter being attached to a female Luer lock tip with hot glue at its non-threaded end. The hot glue should cover the entire attachment point to fully seal the connection. (**C**) The syringe with a 3-way valve attached is connected to the pressure transducer using a 3D-printed adapter. Picture shows the device taken apart before testing to load zebrafish larvae through the 3-way valve into the syringe. (**D**) Wiring diagram of the solenoid and the pressure transducer. The +5 V and GND terminals of the Arduino Uno Rev3 were connected to the breadboard rails, and the pressure transducer’s red, black, and green wires were connected to the positive rail, negative rail, and analog input A1. A TIP120 transistor was connected with a 1 kΩ resistor to pin 13, its middle pin to the solenoid’s GND and a flyback diode 1N4007, and its right pin to the negative rail. The Arduino was then connected to a laptop with the Arduino IDE, and the solenoid was activated when powered. (**E**) An image of the ZePID device.

The pressure transducer and solenoid were wired to a breadboard following the layout in Figure 2D. Power (5 V) and ground rails from the Arduino were connected to the breadboard, and the transducer output (Vout) was routed to analog input A1. A TIP120 transistor served as a switch for the solenoid, driven by Arduino pin 13 through a 1-kΩ resistor; a 1N4007 flyback diode protected the circuit from voltage spikes during solenoid switching. The Arduino program (Supplementary Code) initializes continuous pressure recording and triggers the piston after a preset delay, ensuring synchronized data acquisition and actuation.

To ensure proper mechanical alignment, we designed and 3D-printed a piston holder using PLA filament to secure the pneumatic piston and syringe in place (Figure 2E & Supplementary Figure 1B). The holder prevents lateral movement of the syringe and distributes impact forces more evenly, effectively eliminating syringe breakage, a common issue encountered in weight-drop trials.

For each experiment, 6-dpf larvae were anesthetized in 0.8 mg/mL tricaine and loaded into the syringe chamber by disconnecting the adapter from the 3-way valve. After drawing 10–15 larvae into the syringe, air bubbles were removed and the volume adjusted to 1 mL. The syringe was then resealed, the reservoir was pressurized to the desired level using an external air compressor, and the Arduino was activated to fire the piston and generate a pressure wave through the syringe and across the larval bodies.

### ZePID improves reproducibility of pressure delivery

Pressure measurements were acquired using the integrated pressure transducer, which relayed analog signals to the Arduino and subsequently to a laptop for real-time visualization in the Arduino IDE Serial Monitor. As the piston compressed the syringe chamber, transducer output values increased accordingly. These recordings were exported, and maximum pressures for each trial and condition were extracted and analyzed.

To directly compare ZePID with the traditional weight-drop paradigm, we replicated the weight-drop method using two 100-cm PVC guide tubes of different diameters (large tube: ID 3.418 cm, OD 4.250 cm; small tube: ID 2.600 cm, OD 3.360 cm). Weights greater than 100 g could not be tested because the resulting impacts exceeded syringe durability, causing structural failure.

Weight-drop pressures varied considerably and were strongly affected by guide-tube diameter, despite identical drop heights and masses (Figure 3A). This demonstrates the sensitivity of weight-drop force delivery to minor geometric or frictional differences and the inherent variability of the traditional method. Compared to the traditional weight-drop method, ZePID produced markedly more consistent pressure delivery. Within-group standard deviations were substantially lower for ZePID (Figure 3B) compared to weight-drop (Figure 3A), indicating improved reproducibility under identical operating parameters.

**Figure 3.**
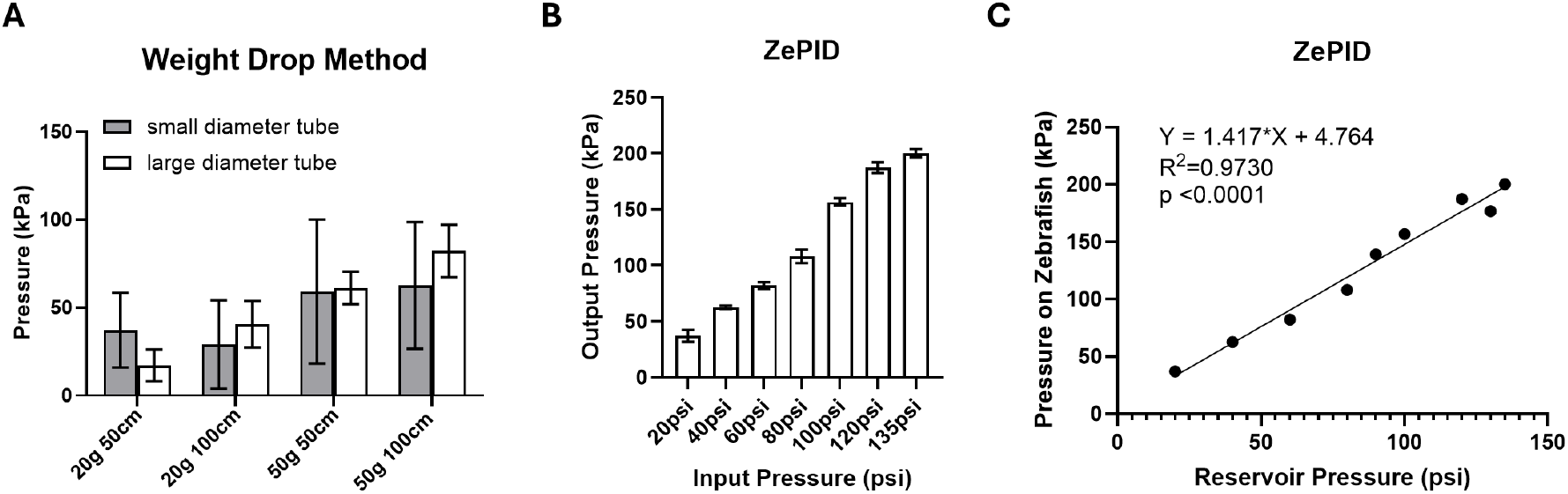
Validating the consistency and predictability of ZePID pressure delivery. (**A**) Measuring maximum pressures inside the syringe achieved by the traditional weight-drop method. Standard deviations show a high level of variation and inconsistency among measurements. Differences were found between large and small tubes despite the same weights being applied. n=5 independent tests. (**B**) Maximum pressure measurements of the ZePID method comparing pressure in the reservoir (Input Pressure) with pressure on zebrafish larvae (Output Pressure). Standard deviations are consistently small among pressures tested. n=5 independent tests. (**C**) Correlation between reservoir pressure and pressure applied to larval zebrafish inside the syringe.

To evaluate the precision of pneumatic pressure modulation, we next plotted the maximum measured syringe pressure against the reservoir pressure used for each trial (Figure 3C). Linear regression revealed a strong positive correlation between reservoir input and delivered pressure (R^2^ = 0.9730, *p* < 0.0001) and the predictive relationship: syringe pressure (kPa) = 1.417× reservoir pressure (psi) + 4.764. This level of control represents a major improvement over the traditional weight-drop approach and provides a quantitative framework for selecting reproducible injury severities.

### Behavioral validation of ZePID-induced traumatic brain injury

To determine whether ZePID-generated pressure pulses produce measurable TBI-associated seizure-like behavioral outcomes^31,32^, we exposed 6-dpf larvae to reservoir pressures of 40, 80, 100, or 150 psi and quantified locomotor behavior 45 minutes post-injury. Individual larvae were placed into 96-well plates and recorded for 10 minutes. Videos were processed in Bonsai^35^ to extract x–y coordinates for each frame, and swimming speeds were averaged across 10-second intervals.

For each fish, the maximum moving speed in 10 second intervals was used as a summary measure and plotted across treatment groups (Figure 4A). One-way ANOVA followed by Dunnett’s post hoc test revealed a significant effect of pressure at 150 psi (*p* = 0.00009), with a clear difference identified between controls (mean ± SD: 56.16 ± 30.82 pixels/s, n = 13) and larvae exposed to pressure (142.43 ± 46.80 pixels/s, n = 8), indicating robust hyperactivity consistent with seizure-like behavior. Lower-pressure groups (40–100 psi) did not differ significantly from controls.

**Figure 4.**
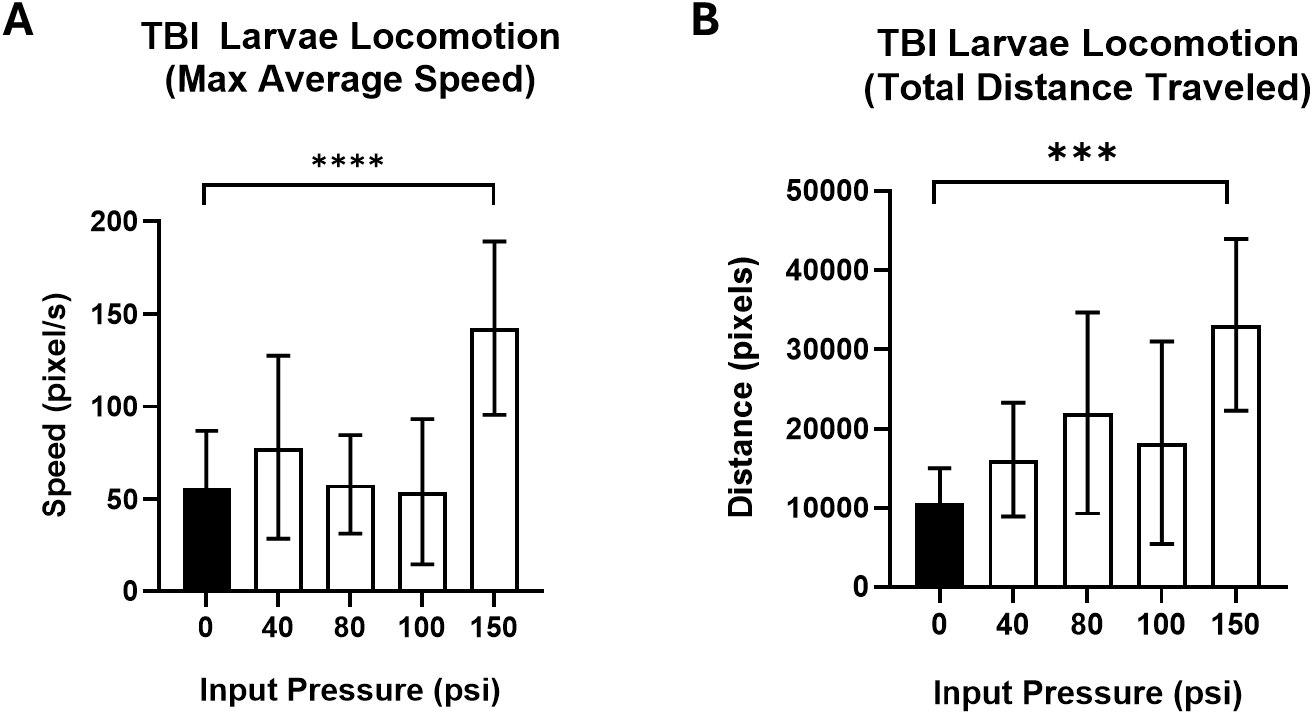
Assessing locomotor behavioral outcomes of ZePID-induced TBI. (**A**) Locomotor assay of larvae following ZePID pressure pulses, showing maximum 10 s interval-averaged speed and standard deviation. Number of larvae per group: 0 psi, n=13; 40 psi, n=4; 80 psi, n=6; 100 psi, n=11; 150 psi, n=8. (**B**) Locomotor assay of larvae following ZePID pressure pulses, showing total distance traveled and standard deviation. Number of larvae per group: 0 psi, n=13; 40 psi, n=4; 80 psi, n=6; 100 psi, n=11; 150 psi, n=8. Significance was calculated by one-way ANOVA and Dunnett’s multiple comparison test. \*\*\**p*<0.001, \*\*\*\**p*<0.0001.

We next quantified the total distance traveled during the 10-minute recording (Figure 4B). Consistent with the speed data, larvae exposed to 150 psi exhibited a significant increase in locomotor activity compared with controls (*p* = 0.0004), traveling 33,134 ± 10,801 pixels (n = 8) versus 10,747 ± 4,297 pixels (n = 13) in the control group. Although the 40- and 80-psi groups showed mild upward trends, these differences were not statistically significant. Together, these behavioral measurements demonstrate that ZePID reliably induces TBI-associated hyperactivity at 150 psi, validating the device as an effective platform for delivering controlled, pressure-based neurotrauma in larval zebrafish.

## DISCUSSION

This study introduces a novel pneumatic system, ZePID, for inducing traumatic brain injury (TBI) in larval zebrafish with substantially greater accuracy, reproducibility, and practicality than the traditional weight-drop method. By replacing the free-fall mass with an electronically controlled pneumatic piston, we achieved highly predictable force delivery, with syringe pressure correlating strongly with reservoir pressure (R^2^ = 0.973). This relationship enables users to precisely select injury severity using the derived calibration equation. In addition, the device footprint was reduced by nearly 70%, making ZePID easier to set up in standard laboratory environment and increasing its accessibility for laboratories with limited bench space.

Behavioral assays demonstrated that ZePID reliably induces TBI-associated hyperactivity at 150 psi, as evidenced by significant increases in both maximum swimming speed and total distance traveled. These seizure-like phenotypes are consistent with established zebrafish TBI models, including weight-drop approaches in larvae^31,32^. The alignment between our results and those of prior weight-drop studies further validates pneumatic delivery as a biologically meaningful and mechanistically comparable injury modality.

Beyond improving reproducibility, ZePID offers flexibility for future applications. Because injury magnitude is determined by reservoir pressure and piston capacity, the device can be readily scaled to accommodate more larvae or older (larger) zebrafish by substituting higher-capacity pneumatic components. This standardization has practical implications for high-throughput drug screening, where consistent injury severity is critical for detecting therapeutic effects without confounding variability introduced during model generation.

Despite the advantages, a limitation of the current device is that at pressures above 150 psi, potential deformation of the device frame may reduce effective force delivery. This necessitates the use of higher-rigidity materials other than PLA filaments in future iterations of this device.

Overall, ZePID provides a precise, scalable, and efficient method for inducing TBI in larval zebrafish and represents a meaningful improvement over the traditional weight-drop method. Future enhancements to the device components and structure combined with broader testing across developmental stages will further expand its utility for mechanistic neurotrauma research and therapeutic discovery.

## MATERIALS AND METHODS

### Zebrafish husbandry

Zebrafish were housed at 26°C–27°C on a 14-hour light, 10-hour dark light cycle. Wild-type AB strain was used for all experiments. All zebrafish experiments were approved by the Institutional Animal Care and Use Committee at the University of Washington.

### Building the ZePID

ZePID consists of two major components: (1) a syringe-based main body that houses the zebrafish larvae and records applied pressure, and (2) a pneumatic actuation system that delivers controlled pressure pulses. These components are connected through Electrical wiring, controlled by Arduino, and held together by a 3D-printed piston holder.

The main body includes a 20-mL Luer-lock syringe, a 3-way Luer valve, and a custom-designed adapter. We used 3-way valves containing two female Luer-lock ports and one male port; the female port opposite the male connector was attached directly to the syringe tip. To interface the valve with the pressure transducer, we designed a cylindrical 3D-printed adapter using Fusion 360 (Supplementary 3D Design 1). The adapter contains internal threads on one side and a narrow, non-threaded projection on the other. A female Luer-lock tip cut from a separate valve using a precision saw was affixed to the projection with hot glue, enabling secure connection to the 3-way valve. The threaded end of the adapter was screwed into a pressure transducer (AUTEX 702373916802), forming a sealed measurement pathway. During experiments, the device is opened at the adapter– valve junction to load larval zebrafish; all other fittings remain permanently assembled to ensure stability and repeatability across trials.

The pneumatic subsystem comprises an air compressor (MOHEGIA), an air reservoir (Vex Robotics), a 2-position, 5-port solenoid valve (SMC SY3120-6LZD-M5), and a 10-mm bore, 10-mm stroke pneumatic piston (Baomain CDJ2B10-10-B). Three 4-mm × M5 push-to-connect fittings were attached to the A, B, and P ports of the solenoid valve. Polyurethane tubing (4-mm OD, 2.5-mm ID) was used to connect 40-cm tubing lengths to ports A and B, and a 10-cm length to port P. Two additional 4-mm × M5 fittings were installed on the air reservoir. One port was connected to the solenoid’s P port via the 10-cm tubing, while the second reservoir port was connected to a 4-mm-to-6-mm pneumatic adapter (ROZESAZZ) using another 10-cm section of 4-mm tubing. This adapter was then joined to a 5-cm piece of 6-mm OD tubing (Beduan), terminating in a Schrader valve (boeray) for charging the reservoir with compressed air.

The pressure transducer and solenoid valve were integrated with an Arduino Uno Rev3 microcontroller via a breadboard. The transducer’s red and black wires were connected to the +5 V and GND rails, respectively, and its Vout line (green wire) was routed to analog input A1. The solenoid was controlled through a TIP120 transistor: the transistor’s left pin was connected to Arduino pin 13 through a 1-kΩ resistor; the middle pin was joined to the solenoid ground and to one terminal of a 1N4007 flyback diode; and the right pin was connected to the breadboard ground. The opposite end of the diode was connected to the positive rail to suppress inductive voltage spikes during switching. Before powering the Arduino, the air reservoir was pressurized by clamping the compressor line to the Schrader valve and maintaining the connection to minimize pressure loss. When the Arduino program was executed, the transducer began recording immediately, and the solenoid triggered piston firing after a programmed delay.

To stabilize the interface between the piston and the syringe plunger, we designed a 3D-printed piston holder (12.9 × 4.5 × 2.2 cm) using Fusion 360 (Supplementary 3D Design 2) and printed using PLA filament. The holder includes rectangular grooves that align and secure both the solenoid and syringe, preventing lateral displacement during piston actuation and improving the durability of the device.

### Inducing Traumatic Brain Injury using ZePID

Wild-type AB zebrafish larvae at 6 days post-fertilization (dpf) were used for all experiments. To standardize injury severity, each larva was exposed to three consecutive pressure pulses at the designated reservoir pressure. Before loading, larvae were anesthetized in 4 μg/mL tricaine (prepared by diluting a 4 mg/mL stock 1:50 in E3 medium) for 5 minutes. The device was opened at the junction between the 3-way valve and the adapter, and 10–20 anesthetized larvae were drawn into the 20-mL syringe. Excess medium was expelled until the final volume reached 1 mL, and the syringe was inverted and gently tapped to remove all air bubbles. The syringe–adapter interface was then reassembled and secured within the ZePID frame. The air reservoir was charged to the specified pressure using the external compressor. The Arduino microcontroller was connected to a power source, and the ZePID program was executed, triggering the piston to fire automatically. After each pulse, the syringe was lightly tapped to redistribute larvae and ensure uniform exposure. This procedure was repeated three times for every experimental condition.

### Locomotor Assay

Following TBI induction, larvae were transferred to fresh E3 medium and allowed to recover in a 28.5°C incubator for 45 minutes. After recovery, individual larvae were placed into separate wells of a 96-well plate using a pipette. For behavioral recording, the plate was positioned beneath a smartphone camera mounted between two support boxes, with a sheet of white printer paper placed underneath to provide a uniform background. The setup was covered with a rigid board to reduce external visual disturbance. Locomotor activity was recorded for 30 minutes, and the middle 10 minutes of each video were extracted for analysis to minimize artifacts caused by human movement near the apparatus. Videos were processed using Bonsai. Each recording was imported, and the software was used to track larvae positions frame-by-frame. The resulting x–y coordinate data for each fish were exported as CSV files for downstream quantification of swimming speed and distance traveled.

### Statistical analysis

Graphs were generated using GraphPad Prism. One-way analysis of variance (ANOVA) analysis was used to analyze data across multiple groups. Dunnett’s multiple comparisons test was used to correct for multiple comparisons. *P* values less than 0.05 were considered significant.

## Supporting information

Supplementary 3D Design 1

Supplementary 3D Design 2

## ACKNOWLEDGEMENTS

We thank the University of Washington Office of Comparative Medicine for providing zebrafish husbandry support. This work was supported by the National Institute of Environmental Health Sciences (NIEHS) of the NIH under the award number R00ES031050. The content is solely the responsibility of the authors and does not necessarily represent the official views of the NIH.

## AUTHOR CONTRIBUTIONS

K.W. designed and conducted the experiments and analyzed data. P.Z. assisted in conducting experiments. Y.G. conceived the study and interpreted the data. K.W. and Y.G. wrote the manuscript. All authors contributed meaningful insights during discussions and reviewed and approved the final version of the manuscript.

## COMPETING INTERESTS

The authors declare that they have no competing interests.

## DATA AND MATERIALS AVAILABILITY

All data needed to evaluate the conclusions in the paper are present in the paper and/or the Supplementary Materials. Arduino code is presented in Supplementary Code.

## Notes

### Competing Interest Statement

The authors have declared no competing interest.

## REFERENCES

1 Naumenko, Y., Yuryshinetz, I., Zabenko, Y. & Pivneva, T. Mild traumatic brain injury as a pathological process. Heliyon 9, e18342 (2023). PMC10372741.

2 Chan, A. et al. Traumatic brain injuries: a neuropsychological review. Front Behav Neurosci 18, 1326115 (2024). PMC11497466.

3 Galgano, M. et al. Traumatic Brain Injury: Current Treatment Strategies and Future Endeavors. Cell Transplant 26, 1118–1130 (2017). PMC5657730.

4 Orr, T. J. et al. Traumatic Brain Injury: A Comprehensive Review of Biomechanics and Molecular Pathophysiology. World Neurosurg 185, 74–88 (2024).

5 Haarbauer-Krupa, J. et al. Epidemiology of Chronic Effects of Traumatic Brain Injury. J Neurotrauma 38, 3235–3247 (2021). PMC9122127.

6 Choi, Y. et al. Elevated Risk of Stroke in Young Adults After Traumatic Brain Injury: A Nationwide Study of 1 Million Individuals. J Am Heart Assoc 13, e033453 (2024). PMC11963958.

7 Freire, M. A. M. et al. Cellular and Molecular Pathophysiology of Traumatic Brain Injury: What Have We Learned So Far? Biology (Basel) 12 (2023). PMC10452099.

8 Greve, M. W. & Zink, B. J. Pathophysiology of traumatic brain injury. Mt Sinai J Med 76, 97–104 (2009).

9 Ng, S. Y. & Lee, A. Y. W. Traumatic Brain Injuries: Pathophysiology and Potential Therapeutic Targets. Front Cell Neurosci 13, 528 (2019). PMC6890857.

10 Vietor, F. I., Sticher, K. & Ashraf, K. H. The Pathophysiology of Traumatic Brain Injuries and the Rationale Behind Creatine Supplementation as a Potential Therapy: A Review. Mo Med 122, 60–66 (2025). PMC11827660.

11 Datta, S. et al. Traumatic brain injury and immunological outcomes: the double-edged killer. Future Sci OA 9, FSO864 (2023). PMC10203904.

12 Maas, A. I. R. et al. Traumatic brain injury: progress and challenges in prevention, clinical care, and research. Lancet Neurol 21, 1004–1060 (2022). PMC10427240.

13 Pilipovic, K., Jankovic, T., Rajic Bumber, J., Belancic, A. & Mrsic-Pelcic, J. Traumatic Brain Injury: Novel Experimental Approaches and Treatment Possibilities. Life (Basel) 15 (2025). PMC12193815.

14 Syzdykbayev, M. et al. A Modern Approach to the Treatment of Traumatic Brain Injury. Medicines (Basel) 11 (2024). PMC11123131.

15 Xiong, Y., Mahmood, A. & Chopp, M. Animal models of traumatic brain injury. Nat Rev Neurosci 14, 128–142 (2013). PMC3951995.

16 Albert-Weissenberger, C. & Siren, A. L. Experimental traumatic brain injury. Exp Transl Stroke Med 2, 16 (2010). PMC2930598.

17 Fesharaki-Zadeh, A. & Datta, D. An overview of preclinical models of traumatic brain injury (TBI): relevance to pathophysiological mechanisms. Front Cell Neurosci 18, 1371213 (2024). PMC11045909.

18 Marklund, N. Rodent Models of Traumatic Brain Injury: Methods and Challenges. Methods Mol Biol 1462, 29–46 (2016).

19 Zhao, Q., Zhang, J., Li, H., Li, H. & Xie, F. Models of traumatic brain injury-highlights and drawbacks. Front Neurol 14, 1151660 (2023). PMC10309005.

20 Padmakumar, S., Kulkarni, P., Ferris, C. F., Bleier, B. S. & Amiji, M. M. Traumatic brain injury and the development of parkinsonism: Understanding pathophysiology, animal models, and therapeutic targets. Biomed Pharmacother 149, 112812 (2022). PMC9050934.

21 Marmarou, A. et al. A new model of diffuse brain injury in rats. Part I: Pathophysiology and biomechanics. J Neurosurg 80, 291–300 (1994).

22 Saxena, B., Bohra, B. & Lad, K. A. Weight-Drop Method for Inducing Closed Head Diffuse Traumatic Brain Injury. Methods Mol Biol 2761, 569–588 (2024).

23 Machado, C. A. et al. Weight-drop model as a valuable tool to study potential neurobiological processes underlying behavioral and cognitive changes secondary to mild traumatic brain injury. J Neuroimmunol 385, 578242 (2023).

24 Allende Labastida, J., Motamedi, M., Wu, P. & Szczesny, B. Protocol for inducing varying TBI severity in a mouse model using a closed-head, weight-drop, impact-induced acceleration mechanism. STAR Protoc 5, 103370 (2024). PMC11736045.

25 Kane, M. J. et al. A mouse model of human repetitive mild traumatic brain injury. J Neurosci Methods 203, 41–49 (2012). PMC3221913.

26 Murashova, L. & Dyachuk, V. Modeling traumatic brain and neural injuries: insights from zebrafish. Front Mol Neurosci 18, 1552885 (2025). PMC11983547.

27 Tikhonova, M. A. et al. A Novel Laser-Based Zebrafish Model for Studying Traumatic Brain Injury and Its Molecular Targets. Pharmaceutics 14 (2022). PMC9416346.

28 Geng, Y. & Peterson, R. T. The zebrafish subcortical social brain as a model for studying social behavior disorders. Dis Model Mech 12 (2019). PMC6737945.

29 Burgess, H. A. & Burton, E. A. A Critical Review of Zebrafish Neurological Disease Models-The Premise: Neuroanatomical, Cellular and Genetic Homology and Experimental Tractability. Oxf Open Neurosci 2, kvac018 (2023). PMC10464506.

30 Siddiqui, S., Siddiqui, H., Riguene, E. & Nomikos, M. Zebrafish: A Versatile and Powerful Model for Biomedical Research. Bioessays 47, e70080 (2025). PMC12632426.

31 Gill, T. et al. Delivering Traumatic Brain Injury to Larval Zebrafish. Methods Mol Biol 2707, 3–22 (2024).

32 Locskai, L. F. et al. A larval zebrafish model of traumatic brain injury: optimizing the dose of neurotrauma for discovery of treatments and aetiology. Biology Open 14 (2025).

33 Main, B. S., Sloley, S. S., Villapol, S., Zapple, D. N. & Burns, M. P. A Mouse Model of Single and Repetitive Mild Traumatic Brain Injury. J Vis Exp (2017). PMC5608469.

34 Mace, B. E. et al. Optimization of a translational murine model of closed-head traumatic brain injury. Neurol Res 46, 304–317 (2024).

35 Lopes, G. et al. Bonsai: an event-based framework for processing and controlling data streams. Front Neuroinform 9, 7 (2015). PMC4389726.

