## Supplementary Code for "High-Precision Pneumatic Induction of Traumatic Brain Injury in Larval Zebrafish": TBI Supplementary Code.ino

```
float difference = 0;
float baselineVoltage;                     
int solenoidPin = 9;  
int timer = 0;
bool pistonFired = false;
void setup() {   
  Serial.begin(9600); //initiate Serial monitor
  pinMode(A1,INPUT); //initiate pressure transducer
  baselineVoltage = getBaselineVoltage();
  pinMode(solenoidPin, OUTPUT); //initiate solenoid;
}

void loop() {
  timer++;
  int sensorValue = analogRead(A1);
  float voltage = sensorValue*(5.0/1023.0);
  //calculate the change in voltage
  if (abs(voltage - baselineVoltage) < 0.005){
    difference = 0;
  }
  else{
    difference = voltage - baselineVoltage;
  }
  float pressure = (difference)*(150/(4.5 - 0.5));
  Serial.println(pressure*6.89);
  //turn on piston
  if (timer == 400 && !pistonFired) {
    digitalWrite(solenoidPin, HIGH);
  }
  if (timer == 405 && !pistonFired) {
    digitalWrite(solenoidPin, LOW);
    pistonFired = true; // prevent future triggers
  }
}  

float getBaselineVoltage() {
  //set the baseline voltage by calculating the average os first 50 data points
  float sum = 0;
  int numReadings = 50;
  for (int i = 0; i < numReadings; i++) {
    sum += analogRead(A1) * (5.0 / 1023.0);
  }
  return sum / numReadings;
}
```
