## Supplementary Materials for "High-Precision Pneumatic Induction of Traumatic Brain Injury in Larval Zebrafish"

**This PDF file includes:**

Supplementary Figure 1

### SUPPLEMENTARY FIGURE

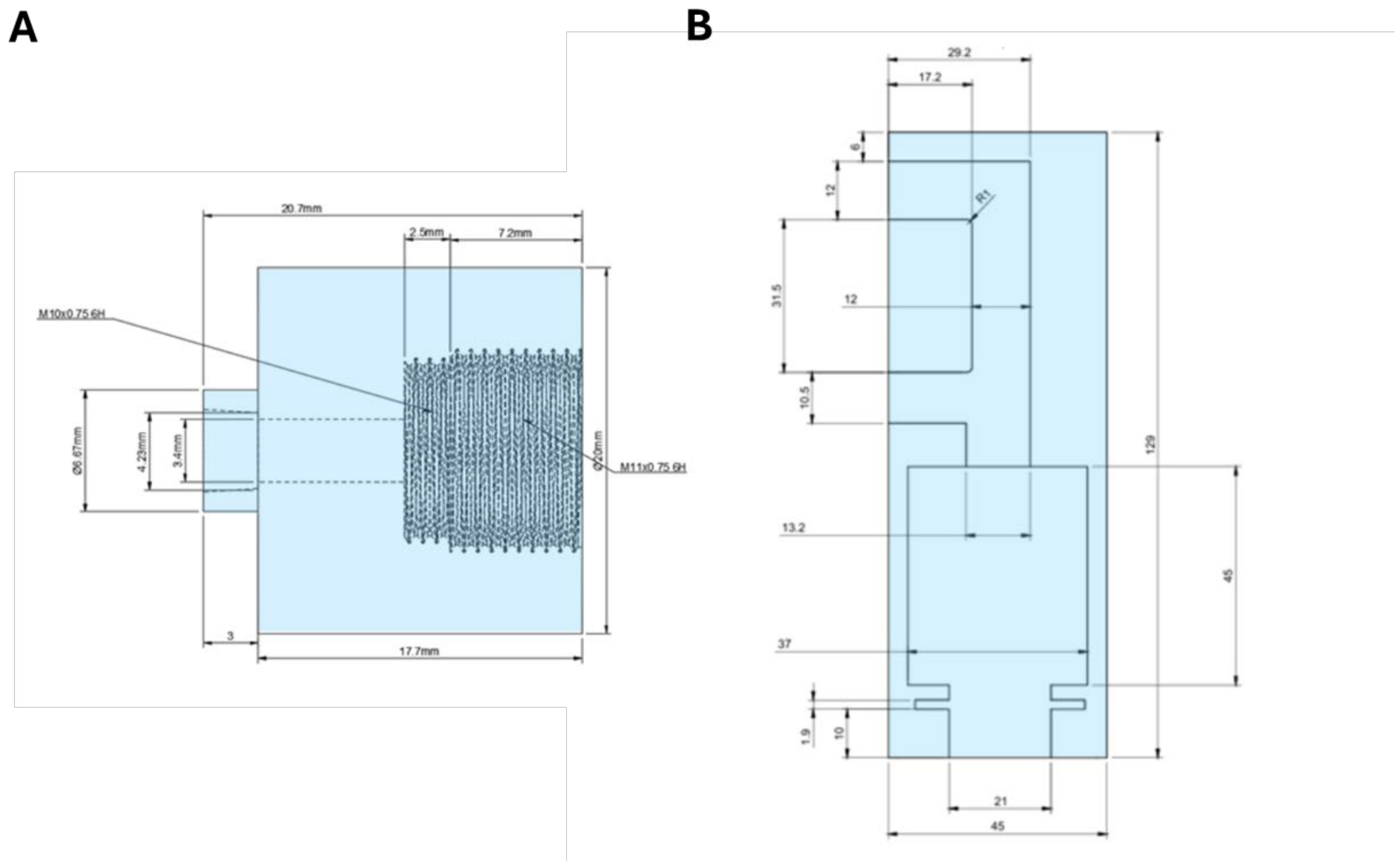


**Supplementary Figure 1.**

(**A**) Design of the custom 3D-printed adapter. (**B**) Design of the 3D-printed piston holder.
